# Polymorphisms in oxidative pathway related genes and susceptibility to inflammatory bowel disease

**DOI:** 10.1101/022681

**Authors:** Nezha Senhaji, Sellama Nadifi, Aurora Serrano, Daniel León Rodríguez, Nadia Serbati, Mehdi Karkouri, Wafaa Badre, Javier Martin

## Abstract

**Objective:** To better characterize the genetic factors, implicated in oxidative pathway, determining susceptibility to inflammatory bowel disease (IBD), we assessed for the first time the potential role of *NOS2A*, *HIF1A* and *NFKB1* polymorphisms on the risk of developing IBD in Moroccan population.

**Methods:** The distribution of (TAAA)n_rs12720460 and (CCTTT)n_rs3833912 *NOS2A* microsatellite repeats, *HIF-1A*_rs11549467 and *NFKB1*–94ins/delATTG_rs28362491 was analyzed in 507 subjects grouped in 199 IBD and 308 healthy controls. Genotyping was performed with polymerase chain reaction-fluorescent method and the TaqMan® allelic discrimination technology.

**Results:** The allele and genotype frequencies of *HIF1A*_rs11549467, *NFKB1*_rs28362491 and *NOS2A*_(TAAA)n did not differ significantly between patients and controls. Analysis of *NOS2A*_(CCTTT)n markers evidenced differences between patients and healthy controls. A preferential presence of the (CCTTT)8 (P=0.02; OR=1.71, 95%CI=1.07–2.74), (CCTTT)14 (P=0.02; OR=1.71, 95%CI=1.06–2.76) alleles in IBD, (CCTTT)8 (P=0.008; OR=1.95, 95%CI=1.17–3.23) in CD and (CCTTT)7 (P=0.009; OR = 7.61, 95%CI=1.25-46.08), (CCTTT)11 (P=0.05 ; OR= 0.51, 95%CI=0.25-1.01), (CCTTT)14 (P=0.02 ; OR= 2.05, 95%CI=1.07-3.94), (CCTTT)15 (P=0.01 ; OR= 2.25, 95%CI=1.16-4.35) repeats in UC patients indicated its possible association with higher disease risk which need to be confirmed in a larger sample size.

**Conclusion:** Our results suggest that the *NOS2A*_(CCTTT)n gene variations may influence IBD susceptibility in the Moroccan population.

## Introduction

Inflammatory bowel disease (IBD), a chronic and relapsing-remitting disorder of the gastrointestinal tract, encompasses Crohn’s disease (CD) and ulcerative colitis (UC).

IBD was thought to be a western disease, with higher prevalence reported in developed countries [1]. However, during the last few decades, incidence rates of the two major forms of IBD have been increasing in developing countries [2], including Morocco.

Chronic intestinal inflammation is a hallmark of both disorders, and is believed to result from a number of abnormal conditions. The involvement of oxidative damage in IBD development has been thoroughly documented. Oxidative stress mainly contributes to aberrant inflammatory responses of intestinal cells to commensal bacteria and dietary antigens. During IBD, activated leukocytes generate a wide spectrum of proinflammatory cytokines, in addition to excessive oxidative reactions that alter the redox equilibrium within the gut mucosa. Therefore, capacity to maintain inflammation by induction of transcription factors and redox-sensitive signaling pathways may influence the occurrence and severity of the disease [3].Induction of inducible nitric oxide synthase (iNOS) was reported to play a key role in oxidative stress-induced inflammation [4]. The genetic polymorphisms of the *NOS2A* (nitric oxide synthase) gene have been proposed to be involved in IBD aetiology[5].Two functionally relevant polymorphisms located at NOS2A gene promoter region were reported, the first one is a highly polymorphic pentanucleotide (CCTTT)n microsatellite repeat which is important in the regulation of NOS2A transcription[6]. The second one is located at the proximal promoter region and consists of an insertion/deletion of one TAAA repeat[7].

Perpetuation of inflammation is also mediated by cellular stress responses of inflammatory cells that produce soluble mediators and reactive species which act by further inducing changes in transcription factors, among which hypoxia-inducible factor-1α (HIF-1α) and nuclear factor κB (NF-κB) [8]. HIF-1-α is a key regulator of cellular response to hypoxia, the gene encoding the HIF-1α subunit (HIF1A) carries a common missense mutation, A588T (G>A, rs11549467), that has been related to increased trans-activation capacity [9]. The involvement of HIF-1-α in the enhancement of the inflammatory response was demonstrated, elevated levels of HIF-1-α in biopsies of primary lesions of patients confirmed its role in inflammatory diseases [10]. HIF-1-α has been shown to induce the secretion of inflammatory mediators by indirect signaling through NF-kB-mediated cytokine and chemokine secretion [11].

NF-κB is activated during inflammation, giving rise to induction of gene expression of several genes involved in mucosal inflammation such as cytokines (TNFA, IL6, IL1β…), Cox-2, and NOS2A [12][13].A functional *NFKB1* promoter polymorphism, consisting of a common insertion/deletion (94ins/delATTG) that seems to affect promoter activity of the *NFKB1* gene and differential nuclear protein binding[14], was associated to the risk of CD [15] and UC [14].

In search of relevant gene polymorphisms related to oxidative stress signaling that are involved in IBD development, we explored the association of *HIF1A*_rs11549467, *NFKB1*_rs28362491 *NOS2A*(CCTTT)n_rs3833912 and *NOS2A*(TAAA)_rs12720460 polymorphisms with IBD (CD and UC) in a Moroccan population.

## Material and methods

### Study population

Peripheral blood was obtained from 311 healthy unrelated blood donors. 199 patients diagnosed with IBD at the CHU Ibn Rochd Hospital (Casablanca, Morocco) were included in this study. The diagnosis of CD or UC was established according to conventional endoscopic, clinical, histological and radiological criteria as previously described[14];[16];[17].CD phenotype was classified according to the Montreal classification [18]. UC anatomic location was subgrouped using Paris classification [19]. Patient’s clinical and demographic characteristics were collected in a case report form including questions on disease location and phenotype, age at diagnosis and other clinical features. The local ethics committee approved the study in accordance with the declaration of Helsinki for experiments involving humans, and a written informed consent was obtained from all participants.

### DNA analysis

Genomic DNA was extracted from peripheral blood using the salting out procedure and from Formalin Fixed Paraffin Embedded Tissues using the QIAamp® DNA FFPE Tissue Kit (Qiagen). DNA quality and quantity was determined with the NanoDrop™ 1000 Spectrophotometer (Thermo Fisher Scientific, Wilmington, DE) and the QuBit Quantification Platform (Invitrogen, Ltd., Paisley, UK) using the QuBit high-sensitivity assay reagents.

The *NOS2A* (TAAA)n_rs12720460 and (CCTTT)n_rs3833912genotyping was performed using a polymerase chain reaction (PCR)-based method combined with fluorescent technology as previously described [20]. Forward and reverse primers were:

F: 5’-TGC CAC TCC GCT CCA G-3’; R: 5’-GGC CTC TGA GAT GTT GGT CTT-3’for (TAAA)n, and F: 5’-ACC CCT GGA AGC CTA CAA CTG CAT-3’; R: 5’-GCC ACT GCACCC TAG CCT GTC TCA-3’ for (CCTTT)n. The forward primers were 5’ labeled with the fluorescent dye 6-Carboxyfluorescein amino hexy FAM. The different alleles were resolved after capillary electrophoresis on automated DNA sequencer (ABI 3130xl Genetic Analyzer, Applied Biosystems) and analyzed with the GeneMapper® 4.0 software (Applied Biosystems). Selected samples from each genotype were sequenced in order to confirm the length of each allele.

Genotyping of *HIF1A* (G/A) rs11549467 and *NFKB1*–94ins/delATTG (rs28362491) was performed on the LightCycler 480 System (Roche, Barcelona, Spain) using a pre-designed TaqMan SNP genotyping assays (Applied Biosystems, Foster City, CA, USA) as previously described [21];[22]. PCR was carried out in a total reaction volume of 5 μl with the following amplification protocol: initial denaturation at 95°C for 3 min followed by 50 cycles of denaturation at 95°C for 3 s, and annealing/extension at 60°C for 20 s. The primer sequences of *NFKB1* promoter polymorphism IN/DEL -94ATTG were: F: 5’-GCC TCC GTG CTG CCT-3’ and R: 5’-AGG GAA GCC CCC AGGAA-3’, and the probe sequences were NFKB1-INS: 5’-VIC-CCCGACCATTGATTGG-NFQ-3’ and NFKB1-DEL: 5’-FAM-TTCCCCGACCATTGG-NFQ-3’.

## Statistical analysis

Genotype and allele distributions among patients with CD, UC and IBD versus healthy controls were compared using the χ^2^ test or Fisher test as appropriate. Odds ratios (ORs) with a confidence interval (CI) of 95% were assessed to measure the strength of association. Statistical power was calculated using Power Calculator of Genetic Studies 2006 software (<u>http://www.sph.umich.edu/csg/abecasis/CaTS/</u>). A chi-square test was used to test for deviation from Hardy-Weinberg equilibrium (HWE). Statistical analyses were performed with Plink software V1.07 (<u>http://pngu.mgh.harvard.edu/purcell/plink/</u>). P-value<0.05 was considered to be statistically significant. Bonferroni correction was applied to significant P-values of *NOS2A* polymorphisms to correct by the number of comparisons.

## Results

Hundred ninety-nine patients with IBD (136 CD; 63 UC) and 308 control subjects were included in this study. The success rates of genotyping assays ranged between 95 and 100%. Baseline demographic and clinical characteristics of cases are presented in a previous report [36].

## *HIF1A* (G/A) rs11549467 and *NFKB1*–94ins/delATTG (rs28362491) polymorphisms

In both patients and controls, the genotype distribution of examined polymorphisms complied with the Hardy-Weinberg expectations.

In order to study associations of *HIF1A* (rs11549467) and *NFKB1* (rs28362491) variants in IBD overall and in CD and UC in particular, the distribution of polymorphic alleles was assessed. Genotype and allele frequencies are given in Tables 1 and 2.

**Table 1:** Minor Allele Frequencies of *HIF1A* and *NFKB1* genetic variants in IBD patients and healthy controls from Morocco.

| Gene | SNP | Group | Number of Alleles | MAF (%) | Allele test |  |
| --- | --- | --- | --- | --- | --- | --- |
|  |  |  |  |  | OR [CI 95%] | P-value |
| *HIF1A* | rs11549467 | Controls (n=308) | 74/542 | 12.01 |  |  |
|  |  | IBD (n=199) | 50/348 | 12.56 | 1.05 [0.72-1.54] | 0.79 |
|  |  | CD (n=136) | 36/236 | 13.24 | 1.12 [0.73-1.71] | 0.61 |
|  |  | UC (n=63) | 14/112 | 11.11 | 0.92 [0.50-1.68] | 0.77 |
| *NFKB1* | rs28362491<br>-94ATTG<br>ins/del | Controls (n=308) | 257/359 | 41.72 |  |  |
|  |  | IBD (n=199) | 167/231 | 41.96 | 1.01 [0.78-1.30] | 0.93 |
|  |  | CD (n=136) | 113/159 | 41.54 | 0.99 [0.74-1.33] | 0.96 |
|  |  | UC (n=63) | 54/72 | 42.86 | 1.05 [0.71-1.54] | 0.81 |

CD: Crohn disease; UC: ulcerative colitis; OR: odds ratio; N: number; MAF: minor allele frequencies

**Table 2:**
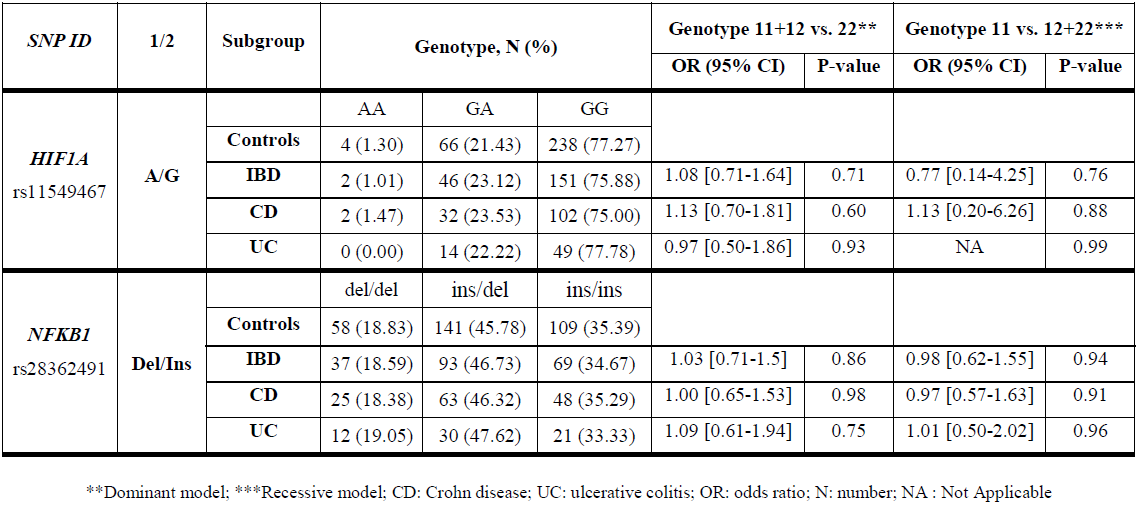
Genotype and genetic models distribution of *HIF1A* and *NFKB1* SNPs in IBD patients and controls.

| SNP ID | 1/2 | Subgroup | Genotype, N (%) |  |  | Genotype 11+12 vs. 22** |  | Genotype 11 vs. 12+22*** |  |
| --- | --- | --- | --- | --- | --- | --- | --- | --- | --- |
|  |  |  |  |  |  | OR (95% CI) | P-value | OR (95% CI) | P-value |
| *HIF1A*<br>rs11549467 | A/G |  | AA | GA | GG |  |  |  |  |
|  |  | Controls | 4 (1.30) | 66 (21.43) | 238 (77.27) |  |  |  |  |
|  |  | IBD | 2 (1.01) | 46 (23.12) | 151 (75.88) | 1.08 [0.71-1.64] | 0.71 | 0.77 [0.14-4.25] | 0.76 |
|  |  | CD | 2 (1.47) | 32 (23.53) | 102 (75.00) | 1.13 [0.70-1.81] | 0.60 | 1.13 [0.20-6.26] | 0.88 |
|  |  | UC | 0 (0.00) | 14 (22.22) | 49 (77.78) | 0.97 [0.50-1.86] | 0.93 | NA | 0.99 |
| *NFKB1*<br>rs28362491 | Del/Ins |  | del/del | ins/del | ins/ins |  |  |  |  |
|  |  | Controls | 58 (18.83) | 141 (45.78) | 109 (35.39) |  |  |  |  |
|  |  | IBD | 37 (18.59) | 93 (46.73) | 69 (34.67) | 1.03 [0.71-1.5] | 0.86 | 0.98 [0.62-1.55] | 0.94 |
|  |  | CD | 25 (18.38) | 63 (46.32) | 48 (35.29) | 1.00 [0.65-1.53] | 0.98 | 0.97 [0.57-1.63] | 0.91 |
|  |  | UC | 12 (19.05) | 30 (47.62) | 21 (33.33) | 1.09 [0.61-1.94] | 0.75 | 1.01 [0.50-2.02] | 0.96 |

\*\*Dominant model; \*\*\*Recessive model; CD: Crohn disease; UC: ulcerative colitis; OR: odds ratio; N: number; NA : Not Applicable

The analysis of both *HIF1A* (rs11549467) and *NFKB1* (rs28362491) polymorphisms distribution among patients and controls did not reveal any statistically significant association, both in terms of allele and genotype frequencies. Similarly, we did not observe any effect on disease risk when Genetic models were assessed (Table 2).

## *NOS2A* (TAAA)n and (CCTTT)n repeat microsatellite polymorphisms

We explored the potential influence of *NOS2A* polymorphisms on the susceptibility to IBD in the Moroccan population. Tables 3 and 4 show the distribution of the pentanucleotide (CCTTT)n microsatellite alleles in controls, IBD, CD and UC cases. Ten different alleles, comprising of 7-16 repeats i.e. 171-216 bp, were observed in our population. (CCTTT)_12_ was observed to be the most frequent allele in IBD (21.2%), CD (21.7%), UC cases (20.0%) and controls (22.0%). The overall (CCTTT)n distribution showed differences between cases and controls.

**Table 3:** Allelic frequencies of (CCTTT)n microsatellite polymorphism of *NOS2A* gene for Moroccan IBD patients and healthy controls.

| Repeat no. | Size (bp) | Controls<br>2n= 596(%) | IBD<br>2n= 382(%) | P value | OR (95%CI) |
| --- | --- | --- | --- | --- | --- |
| 7 | 171 | 2 (0.3) | 4 (1.04) | 0.16 | 3.14 (0.57-17.24) |
| 8 | 176 | 37 (6.2) | 39 (10.2) | **0.02\*** | 1.71 (1.07-2.74) |
| 9 | 181 | 75 (12.5) | 54 (14.1) | 0.48 | 1.14 (0.78-1.66) |
| 10 | 186 | 89 (15.0) | 46 (12.04) | 0.20 | 0.77 (0.53-1.14) |
| 11 | 191 | 90 (15.1) | 45 (11.8) | 0.14 | 0.75 (0.51-1.10) |
| 12 | 196 | 131 (22.0) | 81 (21.2) | 0.77 | 0.95 (0.69-1.30) |
| 13 | 201 | 99 (16.6) | 50 (13.9) | 0.13 | 0.75 (0.52-1.09) |
| 14 | 206 | 36 (6.2) | 38 (9.9) | **0.02\*\*** | 1.71 (1.06-2.76) |
| 15 | 211 | 33 (5.5) | 22 (5.8) | 0.88 | 1.04 (0.59-1.81) |
| 16 | 216 | 4 (0.6) | 3 (0.7) | 0.83 | 1.17 (0.26-5.26) |

\*p= 0.02, OR (95%CI)= 1.71 (1.07-2.74); Bonferroni's corrected $P_c=0.2$ ; \*\*p= 0.02, OR (95%CI)=1.71 (1.06–2.76); $P_c=0.2$

**Table 4:** Allelic frequencies of (CCTTT)n microsatellite polymorphism of *NOS2A* gene for CD, UC patients and healthy controls.

| Repeat no. | Size (bp) | Controls<br>2n= 596(%) | CD<br>2n= 262(%) | P value | OR (95%CI) | UC<br>2n= 120(%) | P value | OR (95%CI) |
| --- | --- | --- | --- | --- | --- | --- | --- | --- |
| 7 | 171 | 2 (0.3) | 1 (0.4) | 0.91 | 1.13 (0.10-12.6) | 3 (2.5) | **0.009<sup>b</sup>** | 7.61 (1.25-46.08) |
| 8 | 176 | 37 (6.2) | 30 (11.4) | **0.008<sup>a</sup>** | 1.95 (1.17-3.23) | 9 (7.5) | 0.59 | 1.22 (0.57-2.61) |
| 9 | 181 | 75 (12.5) | 43 (16.4) | 0.13 | 1.36 (0.90-2.04) | 11 (9.1) | 0.29 | 0.70 (0.36-1.36) |
| 10 | 186 | 89 (15.0) | 33 (12.6) | 0.36 | 0.82 (0.53-1.26) | 13 (10.8) | 0.24 | 0.69 (0.37-1.28) |
| 11 | 191 | 90 (15.1) | 35 (13.3) | 0.50 | 0.86 (0.56-1.32) | 10 (8.3) | **0.05<sup>c</sup>** | 0.51 (0.25-1.01) |
| 12 | 196 | 131 (22.0) | 57 (21.7) | 0.94 | 0.98 (0.69-1.40) | 24 (20.0) | 0.63 | 0.88 (0.54-1.44) |
| 13 | 201 | 99 (16.6) | 31 (11.8) | 0.07 | 0.67 (0.43-1.03) | 19 (15.8) | 0.83 | 0.94 (0.55-1.61) |
| 14 | 206 | 36 (6.2) | 24 (9.1) | 0.09 | 1.56 (0.91-2.68) | 14 (11.6) | **0.02<sup>d</sup>** | 2.05 (1.07-3.94) |
| 15 | 211 | 33 (5.5) | 8 (3.0) | 0.11 | 0.53 (0.24-1.18) | 14 (11.6) | **0.01<sup>e</sup>** | 2.25 (1.16-4.35) |
| 16 | 216 | 4 (0.6) | 0 (0) | 0.18 | 0 (0) | 3 (2.5) | 0.06 | 3.79 (0.83-17.18) |

<sup>a</sup>p=0.008, OR (95%CI)= 1.95 (1.17-3.23); $P_c= 0.08$ ; <sup>b</sup>p=0.009; OR (95%CI)= 7.61 (1.25-46.08); $P_c= 0.09$ ; OR (95%CI)= 0.51 (0.25-1.01); $P_c= 0.5$ ; <sup>d</sup>p=0.02; OR (95%CI)= 2.05 (1.07-3.94); $P_c= 0.2$ ; <sup>e</sup>p=0.01; OR (95%CI)= 2.25 (1.16-4.35)

When individual CCTTT alleles were analyzed, a significant increase in frequency of the 8-repeat (10.2% vs. 6.2%; P=0.02, OR=1.71, 95%CI=1.07–2.74) and 14-repeat (9.9% vs. 6.2%; P=0.02, OR=1.71, 95%CI=1.06–2.76) alleles was observed in IBD cases compared with controls respectively.

Similarly the (CCTTT)8 repeat/allele was found to be higher in CD patients cohort as compared to controls (11.4% vs. 6.2%; P=0.008, OR=1.95, 95%CI=1.17–3.23). Whereas the increased distribution of the (CCTTT)_14_ repeatamong CD cases compared to controls did not reach the significance level (9.1% vs. 6.2%; P=0.09, OR=1.56, 95%CI=0.91–2.68).

Furthermore, determination of allele frequencies in UC patients revealed significant association of the 7, 11, 14 and 15 repeats polymorphic forms of the microsatellite to disease risk. Similar trend, though non-significant, was observed for the 16 repeat (P=0.06, OR=3.79, 95%CI=0.83–17.18).

However, it should be noted that after Bonferroni correction by the number of comparisons, none of the observed associations remained significant in all patient groups.

We further analyzed the distribution of TAAA insertion/deletion polymorphism of *NOS2A* gene in our population. In this regard, no statistically significant allele or genotype differences were observed between IBD patients and controls (Table 5). Nor were significant differences between stratified CD and UC patients when compared to controls.

**Table 5:** Allele and genotype frequencies of *NOS2A* TAAA polymorphism in IBD patients and controls

|  | Controls<br>n= 295 (%) | IBD patients<br>n= 190 (%) | CD patients<br>n= 132 (%) | UC patients<br>n= 58 (%) |
| --- | --- | --- | --- | --- |
| **Genotype** |  |  |  |  |
| 220/220 | 189 (64.07) | 132 (69.47) | 90 (68.18) | 42 (72.41) |
| 220/224 | 97 (32.88) | 49 (25.79) | 34 (25.76) | 15 (25.86) |
| 224/224 | 9 (3.05) | 9 (4.74) | 8 (6.06) | 1 (1.72) |
| **Allele** | **2n= 590** | **2n=380** | **2n=264** | **2n=116** |
| 220 | 475 (80.5) | 313 (82.4) | 214 (81) | 99 (85.3) |
| 224 | 115 (19.5) | 67 (17.6) | 50 (19) | 17 (14.7) |

No statistically significant differences were found in any of the comparisons.

## Discussion

IBD is an inflammatory disease resulting from a compound effect of a number of abnormal conditions. Genetic factors may play a pivotal role in the development of IBD. In this regard, we explored the potential contribution of genetic polymorphisms of genes related to oxidative pathway to the risk of IBD development. Our attention was focused on functionally relevant polymorphisms located in the *NOS2A* and *NFKB1*genes, and a common missense mutation of *HIF1A* gene.

We found differential distribution of (CCTTT)n microsatellite repeats between patients and controls; namely the (CCTTT)8 and (CCTTT)_14_ repeats for IBD, the (CCTTT)8 for CD and the (CCTTT)7, (CCTTT)_11_, (CCTTT)_14_ and (CCTTT)_15_ repeats for UC patients. As for Caucasians, the most common allele observed in our population was the 12 repeats instead of 10 and 11 repeats for Northwestern Colombians [20].To our knowledge only two reports investigated the involvement of *NOS2A* gene polymorphisms in IBD etiology. Concordantly to our results, M C. Martín *et al*., evidenced the influence of the inducible nitric oxide synthase (CCTTT)n microsatellite repeats on UC risk [5]. However, in contrast to our finding J. Oliver *et al*., demonstrated no tendency toward an association with IBD predisposition [23]. Polymorphisms of *NOS2A* have also been involved in other autoimmune diseases such multiple sclerosis and rheumatoid arthritis [24], [25].

In terms of functional relevance, CCTTT polymorphic markers have been described to affect nitric oxide synthase (NOS) transcription [6]. The involvement of the inducible (calcium-independent) isoform, iNOS in inflammation has been largely demonstrated [26] and is directly related to the large amounts of NO produced by the enzyme after transcriptional induction and the injurious levels of RNS generated by activated leukocytes, macrophages and epithelial cells in the intestinal mucosa [27]. The overexpression of iNOS during active IBD is characterized by elevated rectal NO levels [28].Biopsies of UC-active patients demonstrate higher iNOS transcripts and enzyme levels as compared to controls or healthy relatives [29]. It was also demonstrated that in UC, greatly increased production of iNOS-derived NO reacts with tyrosine leading to production of nitrotyrosine which is associated with infiltration of neutrophils in the epithelium [30][31].

In the other hand, the present study sought to assess the association of *HIF1A* (G/A) rs11549467 and *NFKB1*–94ins/delATTG (rs28362491) polymorphisms with IBD among Moroccan patients. Data on association of these genes with IBD in the North African population are currently lacking. Our results suggest that the studied *NFKB1* gene variation do not influence susceptibility to IBD (CD and UC) in the cohort tested herein. Our findings are in accordance with previous investigations analyzing Spanish [32] British [33]and German populations [34]. In contrast, an association of the –94ins/delATTG polymorphism with UC was demonstrated in a North American population [14]. These discrepant results may have been caused by clinical, population and genetic differences in addition to ethnic origin heterogeneity.

In addition, the present case-control study could not establish a role for *HIF1A* (G/A) rs11549467 polymorphism in the pathogenesis of both CD and UC and also found no evidence for disease risk when evaluating genetic models. This later polymorphism was shown to be associated with autoimmune diseases such as systemic sclerosis [35], however no investigation has assessed its involvement in IBD etiology.

## Conclusion

Our results suggest although not formally demonstrate that, variation in the distribution of CCTTT repeats in the *NOS2A* gene may contribute to IBD development in the Moroccan population. Additionally, our data do not support a role for the *NFKB1*and *HIF1A* polymorphisms in the pathogenesis of IBD.

